# Ubiquitin ligase LNX1 is a major regulator of glycine recapture by the presynaptic transporter GlyT2

**DOI:** 10.1101/233213

**Authors:** E Núñez, E Arribas-González, B López-Corcuera, C Aragón, J de Juan-Sanz

**Author notes:** Both authors contributed equally to this work (E Núñez, E Arribas-González). Corresponding Author’s (de Juan-Sanz J), (Aragón C).

## Abstract

The neuronal glycine transporter GlyT2 is an essential regulator of glycinergic neurotransmission that recaptures glycine in presynaptic terminals to facilitate quantal transmitter packaging in synaptic vesicles. Alterations in GlyT2 expression or activity result in lower cytosolic glycine levels, emptying glycinergic synaptic vesicles and impairing neurotransmission. Lack of glycinergic neurotransmission caused by GlyT2 loss-of-function mutations results in Hyperekplexia, a rare neurological disease characterized by generalized stiffness and motor alterations that may result in sudden infant death. Although the importance of GlyT2 in pathology is known, how this transporter is regulated at the molecular level is poorly understood, limiting current therapeutic strategies. Guided by an unbiased screening, we discovered that the E3 ubiquitin ligase Ligand of Numb protein X1 (LNX1) modulates the ubiquitination status of GlyT2. LNX1 ubiquitinates a cytoplasmic C-terminal lysine cluster in GlyT2 (K751, K773, K787 and K791) through its N-terminal RING-finger domain, and this process regulates the expression levels and transport activity of GlyT2 in neurons. These experiments reveal for the first time the identity of an E3 ubiquitin-ligase acting on GlyT2 and identify a novel regulatory mechanism by which neurons regulate GlyT2 expression and activity.

## INTRODUCTION

Glycine acts as an inhibitory neurotransmitter in the central nervous system (CNS), playing a fundamental role in neuronal circuits of the central auditory pathway, receptive fields in the retina and spinal cord sensitive pathways. Glycinergic neurotransmission strength is controlled presynaptically by the activity of a surface glycine transporter, GlyT2, which recaptures glycine back to the presynaptic terminal to refill synaptic vesicles. Alterations in GlyT2 expression or activity result in the emptying of synaptic vesicles, which vastly weakens glycinergic neurotransmission^1,2^. In humans, this dysfunction is the main presynaptic cause of Hyperekplexia^3–6^, but may also result in chronic pain^7^ and deficits in auditory processing^8^. Although the importance of GlyT2-mediated glycine transport in pathology is known^2,6,9^, our knowledge on how the activity of this transporter is regulated is currently limited. Understanding the molecular regulation of GlyT2 transport would provide insight into the molecular and cellular basis of glycinergic neurotransmission and potentially lead to identifying new therapeutic targets for presynaptic Hyperekplexia. Previous studies on GlyT2 regulatory mechanisms revealed that GlyT2 activity is regulated by PKC activation^10,11^, calnexin function^12,13^, P2Y and P2X purinergic receptors^14,15^ and interaction with syntaxin1^16^, Na+/K+-ATPase^17^ and PMCAs^18^. In addition, we previously described that GlyT2 trafficking and surface expression are regulated by ubiquitination^11,19^, a process in which the small protein ubiquitin is covalently attached to a cytoplasmic lysine residue of a target protein. Protein ubiquitination is a versatile regulatory post-translational modification that controls intracellular signaling events essential for neuronal function and synapse integrity, including trafficking and turnover of presynaptic proteins^20–22^. We discovered that ubiquitination of a cytoplasmic C-terminal lysine cluster in GlyT2 (K751, K773, K787 and K791) controls GlyT2 activity by regulating surface and total expression of the transporter^19^. However, the molecular identity of the enzymes catalyzing GlyT2 ubiquitination remains unknown.

The enzymatic cascade catalyzing ubiquitination of any substrate comprises the sequential activity of the E1 ubiquitin-activating enzyme, E2 ubiquitin-conjugating enzyme and E3 ubiquitin-ligase, which provides specificity to the reaction^23^. E3s have dual roles as both molecular matchmakers and catalysts, bringing together the right E2 with the proper substrate and greatly increasing the rate of ubiquitin transfer^24,25^. Ligand of Numb protein X1 (LNX1) is a RING-type E3 ubiquitin ligase that was first identified as an interacting partner for Numb, a cell fate determinant protein that promotes neuronal differentiation and maturation during the nervous system development^26,27^. LNX1 presents two main isoforms, p70 and p80, with p70 not containing a RING-finger domain. LNX1 is mainly expressed in the nervous system^28^ and presents four consecutive PDZ domains in its structure, which promote its interaction with many other neuronal substrates such as c-Src^29^, PKCα^30^ or the presynaptic active zone proteins CAST^31^, ERC1, ERC2 and LIPRIN-αs^32^. Previous efforts to identify potential LNX1 substrates have used proteomic approaches to identify PDZ binding partners for each of the PDZ domains in LNX1^33,34^. In particular, the second PDZ domain of LNX1 (PDZ2) is a class I PDZ domain, which binds C-terminal motifs with the sequence S/T-X-C^33^. The screening for LNX1 PDZ2 domain reported 26 candidate binding partners, including GlyT2, which contains a PDZ binding motif in its C-terminus with sequence TQC^33^. Given the importance of GlyT2 in the control of inhibitory glycinergic neurotransmission and that ubiquitination is an essential post-translational modification that regulates its function and expression, we decided to explore whether LNX1 is a presynaptic E3-ligase that controls GlyT2 activity.

In this work we demonstrate that GlyT2 interacts with and it is ubiquitinated by LNX1. Increased LNX1 expression reduces total and surface expression of GlyT2, which impairs glycine transport in neurons. However, when LNX1 E3 ligase activity is inactivated by a point mutation in the RING-finger domain, LNX1 is unable to regulate GlyT2 expression and transport activity. In addition, mutations in the C-terminal lysine cluster of GlyT2 prevent both ubiquitination and effects on GlyT2 activity induced by wild type LNX1, indicating that the last 4 lysines are necessary for GlyT2 ubiquitination and thus, control its function and expression levels. Taken together, these findings indicate that LNX1 activity can modulate presynaptic glycine recapture by regulating GlyT2 ubiquitination levels and expression, suggesting that LNX1 may play a role in modulating inhibitory glycinergic neurotransmission strength in the CNS.

## EXPERIMENTAL PROCEDURES

### Materials

Male Wistar rats were bred under standard conditions at the Centro de Biología Molecular Severo Ochoa in accordance with the current guidelines for the use of animals in research. All animal procedures were approved by the institutional animal care and performed according to European Union guidelines (Council Directive 2010/63/EU). Antibodies against GlyT2 n-terminus and GST were generated in house (rabbit and rat: ^35,36^) while the other primary antibodies used were: anti-c-Myc (DHSB, clon 9E10), anti-FLAG M2 (Sigma-Aldrich, F3165), anti-HA (Sigma-Aldrich, clon 12CA5), anti-LNX1 (Novus Biologicals, NBP1-80518), anti-α-tubulin (Sigma-Aldrich, T6074). All chemicals used were from Sigma Aldrich unless otherwise noticed. Neurobasal medium and B27 supplement were purchased from Invitrogen.

### Plasmid constructs

FLAG-tagged p80-LNX1 and p80-LNX1^C45A^ were a generous gift from Prof. Jane McGlade (University of Toronto), Myc-tagged p70-LNX1 was a gift from Yutaka Hata (Addgene plasmid #37009) and pRK5-HA-Ubiquitin-WT was a gift from Ted Dawson (Addgene plasmid #17608). Generation of GFP-GlyT2 and GlyT2-4KR was described in previous work^16,19^. GlyT2-Δ8cter was generated by introducing a stop codon at the residue D792 by site-directed mutagenesis as previously described^37^. GST fused to the C-terminus of GlyT2 was generated by fusion of a 208 bp fragment of GlyT2 cDNA 3’region encoding the C-terminal 68 amino acids of the protein to GST using the PGEX5X vector. The fusion protein was purified from bacterial lysates after IPTG induction. All GlyT2 constructs are based on the longest rattus novergicus isoform of 799 aminoacids (Slc6a5-201, Ensembl transcript ID ENSRN0T00000041950.4). p80-LNX1 and p80-LNX1^C45A^ were cloned in pmCherry-C1 (Clontech) between EcoRI and BamHI sites, generating constructs that express mCherry followed by a linker (SGLRSRAQASNS) and the sequence of each p80-LNX1 variant.

### Cell Growth and Protein Expression

—COS 7 cells (American Type Culture Collection) were grown at 37 °C and 5% CO2 in Dulbecco’s modified Eagle’s medium (DMEM) supplemented with 10% fetal bovine serum. Transient expression was achieved using TrueFect^™^ (United Biosystems), according to the manufacturer’s protocol, and cells were then incubated for 48 h at 37 °C. Reproducible results were obtained with 80–90% confluent cells on 60-mm or 6-well plates, using 5 and 2 μg of total DNA, respectively. The ratio of DNA between GlyT2 and LNX1 variants was 1:4 to favor LNX1 overexpression.

### Primary Cultures of Cerebral Cortex and Transfection

—Primary cultures of embryonic cortical neurons were prepared as described previously^12^. Briefly, the cortex of Wistar rat fetuses was obtained on the 18th day of gestation, and the tissue was mechanically disaggregated in Hanks’ balanced salt solution (Invitrogen) containing 0.25% trypsin (Invitrogen) and 4 mg/ml DNase (Sigma). Cells were plated at a density of 500,000 cells/well in 6-well plates (Falcon), and they were incubated for 4 h in DMEM 10% FCS, containing glucose (10 mM), sodium pyruvate (10 mM), glutamine (0.5 mM), gentamicin (0.05 mg/ml) and streptomycin (0.1 mg/ml). After 4 h, the buffer was replaced with Neurobasal/B27 culture medium containing glutamine (0.5 mM, 50:1 by volume; Invitrogen), and 3 days later, cytosine arabinoside (2μM) was added to inhibit further glial growth. For transfection, neurons that had been maintained *in vitro* for 7 days were incubated with 2μg of total DNA mixed with 4μl of Lipofectamine 2000 reagent (Invitrogen). The ratio of DNA between GlyT2 and LNX1 was 1:3 for both cell imaging and glycine transport experiments to favor LNX1 overexpression. Expression of GlyT2 and LNX1 constructs was always assayed 24h post-transfection in neurons (DIV8), as LNX1 presents a very short half-life and its expression is lost after 48h in primary neurons.

### Immunofluorescence microscopy

Live-cell imaging was performed using a custom-built laser illuminated epifluorescence microscope with an Andor iXon+ (model #DU-897E-BV) back-illuminated electron-multiplying charge-coupled device camera as described previously^38^. Fluorescence imaging was performed by illuminating cells with a 488 or 532 nm lasers and all images were acquired through a 40 × 1.3 NA Zeiss objective. Laser power was constant between experiments and was ~0.35 mW at the back aperture.

### Immunoprecipitation and western blotting

COS7 cells were lysed for 30 min at room temperature (RT) at a concentration of 1.5 mg of protein/ml in TN buffer (25 mM Tris HCl and 150 mM NaCl [pH 7.4]) containing 0.25% Nonidet P-40 (NP-40) and protease inhibitors (PIs: 0.4 mM phenylmethylsulfonyl fluoride (PMSF) and Sigma cocktail). After 15 min centrifugation in a microfuge to remove the cell debris, 4 % of protein was separated to quantify total protein (T) and 5 μl of the primary antibody were added and left overnight at 4°C using the following antibodies for immunoprecipitation: rat anti-GlyT2, anti-c-Myc or anti-FLAG M2. A negative control was also run in parallel in which no antibody was added. Subsequently, 50 μl of 50% protein G agarose beads (PGA; ABT beads inc.) were added and incubated for 45 min at 4°C. The beads were collected by mild centrifugation and washed twice for 7 minutes with lysis buffer at RT. Finally, the beads were pelleted and the immunoprecipitated proteins were eluted in Laemmli buffer at 75°C for 10 min, resolved in SDS/PAGE gels (7.5%), detected in Western blots by enhanced chemiluminescence (ECL), imaged on a GS-810 imaging densitometer (Bio-Rad) and quantified using ImageJ.

### Glutathione S-Transferase Pull-Down Assay

Experiments were performed as previously described^39^. Briefly, COS7 cells transfected with FLAG-tagged p80-LNX1 were lysed in 1 mL of the NP40 lysis buffer at 4°C for 30 min. Samples were centrifuged at 15.000 rpm at 4°C for 10 min to collect the supernatant. After precleaning the cell lysate with GST-glutathione-agarose beads, equal amounts of protein were incubated with 40 μl of GST fusion proteins (GST fused to the N-terminus of GlyT2^35^ or GST fused to the C-terminus of GlyT2), immobilized on glutathione agarose beads for 2 h at 4°C end-over-end mixing. Beads were washed four times with 1 ml of ice-cold lysis buffer. The bound proteins were eluted by adding 50 μl of 20 mM reduced glutathione in 50 mM Tris-ClH (pH 8.0). Samples were analyzed by western blot with anti-FLAG antibody and GST fusion protein expression was confirmed using an anti-GST antibody.

### Ubiquitination assays

COS7 cells were transiently transfected with plasmids encoding wild-type or mutant GlyT2 proteins and HA-tagged ubiquitin. 48 hours after transfection, cells were treated for 3 hours at 37°C with 10 μM MG132. Cells were washed twice with PBS at 4°C, harvested in 50mM Tris, 150mM NaCl, 50mM N-ethylmaleimide buffer with protease inhibitors PMSF and Sigma protease inhibitor cocktail (Ubiquitination Buffer, UB) and analyzed for protein quantification. Equal amounts of protein samples were centrifuged and pellets was resuspended in 90 microliters of UB. 10 microliters of 10% sodium dodecyl sulphate (SDS) were added and incubated for 10 min at 95°C to eliminate protein interactions. Cell lysates were collected and immunoprecipitated with the indicated antibodies and immunoblotted, as described.

### Surface Biotinylation

COS7 cells expressing wild type or mutant GlyT2 were grown in 6-well plates, washed in cold PBS, and labeled with Sulfo-NHS-Biotin (1.0 mg/ml in PBS; Pierce) at 4 °C, a temperature that blocks membrane trafficking of proteins. After quenching with 100 mM L-lysine to inactivate free biotin, the cells were lysed with 1 ml lysis buffer as described elsewhere^13^. A portion of the lysate was saved to determine the total protein content (shown in figures as T), and the remainder was incubated with streptavidin-agarose beads for 2h at room temperature with constant shaking. After centrifugation, the supernatant was removed. The agarose beads recovered were washed three times with 1ml lysis buffer and bound proteins (biotinylated) were eluted with Laemmli buffer (65 mM Tris, 10% glycerol, 2.3% SDS, 100 mM DTT, 0.01% bromphenol blue) for 10 min at 75 °C. The samples were then analyzed in Western blots, where biotinylated samples are noted as BT.

### [^3^H]-Glycine transport assays

For GlyT2 activity determination, the uptake solution contained an isotopic dilution containing 2 μCi/ml [^3^H]glycine (1.6 TBq/mmol; PerkinElmer Life Sciences) in PBS yielding a 1μM or 10μM final glycine concentration in COS7 cells or transfected cortical neurons, respectively, in the absence or presence of the GlyT2 antagonist ALX1393 (0.4 μM,IC50 = 50 nM) to measure background glycine accumulation. Transport was measured by subtracting the background glycine accumulation in COS7 cells or primary neurons and normalizing to the protein concentration. Experiments in neurons were performed in the presence of a GlyT1 antagonist (10 μM NFPS) as this cultures present GlyT1 endogenous activity. The reactions were terminated after 10 min by aspiration followed by washing with HBS. Transport experiments in neurons are performed in primary cortical neurons since they do not express GlyT2 endogenously, which allows expressing GlyT2 mutants and measure their activity without having wild type GlyT2 background transport. Additionally, given the low efficiency of transfection in primary cultures of neurons (~5%), this facilitates measuring glycine transport specifically in GlyT2/LNX1 co-transfected neurons. Similar experiments in spinal cord neurons result in a dilution of the effect since ~95% of the neurons endogenously expressing GlyT2 do not overexpress LNX1 (not shown), making it difficult to dissect the effects of LNX1.

### Densitometry and Data Analysis

The protein bands visualized by ECL (Amersham Biosciences) or fluorography were imaged in a GS-800 calibrated imaging densitometer and quantified using ImageJ, with film exposures in the linear range. All statistical analyses were performed using Origin 8.0 (OriginLab Corp, MA). One-way analysis of variance (ANOVA) was used to compare multiple groups, with subsequent Tukey’s post-hoc test to determine the significant differences between samples. The Student’s *t*-test was used to compare two separate groups. p values are denoted through the text as follows: *, p<0.05; **, p<0.01; ***, p<0.001; p<0.05 or lower values were considered significantly different when compared by one-way ANOVA (Tukey’s posthoc test) or Student’s *t*-test. Thorough the text, the Box whisker plots represent median (line), mean (point), 25 – 75 percentile (box), 10 – 90 percentile (whisker), 1 – 99 percentile (X) and min – max (–) ranges.

## RESULTS

### Ubiquitin E3 ligase LNX1 interacts with GlyT2

GlyT2 surface expression is strongly regulated by ubiquitination but the identity of the E3 ligase that catalyzes this control mechanism remains elusive. The last eight amino acids of GlyT2 were found to bind the PDZ2 domain of E3 ligase LNX1 in a random peptide screening (Fig. 1A). This binding site in GlyT2, which contains a C-terminal PDZ binding motif (PBM), is evolutionarily conserved in most animal species (as evident in sequence alignments: Fig. 1B), suggesting an important role of this sequence for GlyT2 function. To confirm the predicted interaction of LNX1 with the C-terminus of GlyT2^33^, we performed a pulldown assay using three different baits: purified GST, GST fused to the C-terminus of GlyT2 or GST fused to the N-terminus of GlyT2, and combined these with a cell lysate of COS7 cells expressing FLAG-tagged LNX1 isoform p80 (p80-LNX1), the longest LNX1 isoform that contains a functional RING finger domain responsible for the ubiquitin-ligase activity (Fig 1A). We used FLAG immunoblotting to the detect LNX1 co-purification in these experiments and observed that while no LNX1 was found co-purified in either the GST control or the condition using GST-N-terminus, a clear interaction was observed when the GlyT2 GST-C-terminus fusion protein was used (Fig 1C), confirming that LNX1 interacts with GlyT2 through its C-terminal sequence. Immunoblotting anti-GST confirmed the amount of fusion protein used, showing different sizes corresponding to GST fused to the C-terminus (68 amino acids) and N-terminus (193 amino acids) of GlyT2. Next, to explore whether LNX1 could be an E3 ligase for GlyT2, we tested whether the two full-length proteins interact in heterologous cells. We transfected COS7 cells with GlyT2 and FLAG-tagged p80-LNX1 and used coimmunoprecipitation assays to detect whether these proteins interact. Immunoprecipitation of GlyT2 resulted in coimmunoprecipitation of FLAG-tagged p80-LNX1 (Fig 1D, second lane), and correspondingly, immunoprecipitation of FLAG-tagged p80-LNX1 coimmunoprecipitated GlyT2 (Fig 1D, third lane). We also explored whether p70-LNX1, a shorter isoform with no E3 ligase activity (Fig 1A), interacts with GlyT2, as this isoform also contains the PDZ II domain that was predicted to bind the transporter. Using the same approach, we observed that immunoprecipitation of GlyT2 resulted in coimmunoprecipitation of myc-tagged p70-LNX1 (Fig 1E, second lane), and correspondingly, immunoprecipitation of myc-tagged p70-LNX1 coimmunoprecipitated GlyT2 (Fig 1E, third lane; see high exposure). These signals are specific as they are only observed in cells expressing both proteins (data not shown), indicating that the previously predicted interaction between the PDZ II domain of LNX1 and the last 8 amino acids of GlyT2^33^ occurs between both full-length proteins in a cellular environment. Additionally, given that both GlyT2 and LNX1 are expressed in spinal cord neurons^28,35^, we tested whether this interaction also occurs in a neuronal environment following a similar strategy using immunoprecipitation. These experiments were performed in primary cortical neurons since they do not express GlyT2 endogenously, which allows controlling for specificity of the interaction as one can identify nonspecific artifacts in cortical neurons expressing only p80-LNX1 (Fig 1F, left). Immunoprecipitation of GlyT2 also coimmunoprecipitated FLAG-tagged p80-LNX1 in neurons (Fig 1F, right, second lane), confirming the initial results obtained in COS7 cells and indicating that this interaction occurs in a neuronal environment where both proteins are endogenously expressed^28,35^.

**Figure 1.**
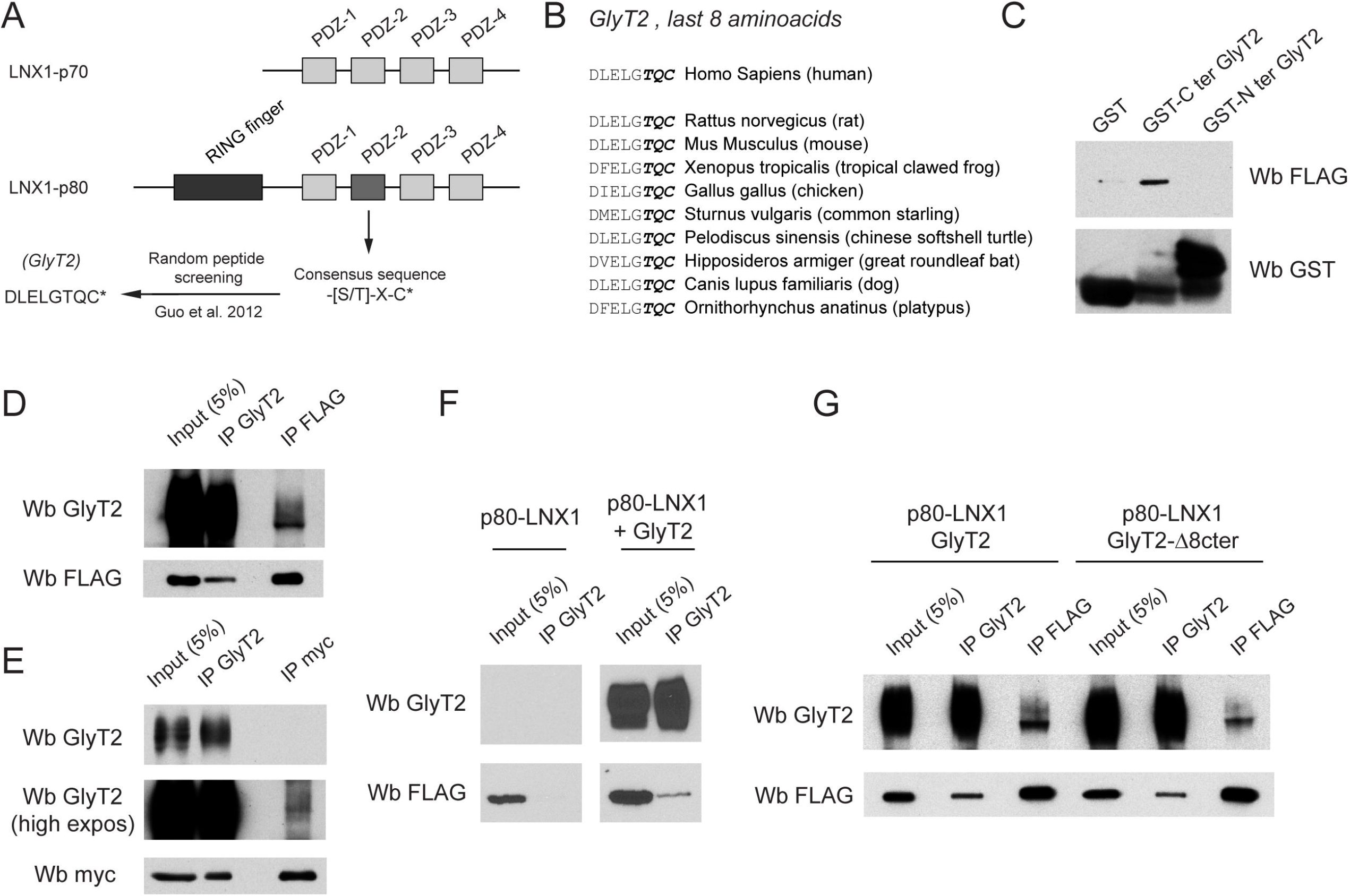
GlyT2 interacts with LNX1 in heterologous cells and primary neurons. A) Graphic scheme showing the two main isoforms of LNX1, p70 (short isoform) and p80 (long isoform with RING domain). A previous unbiased screening effort by Guo et al^33^ identified the last 8 amino acids of GlyT2 (DLELGTQC) as a possible interacting peptide for the PDZ II domain of LNX1, which recognizes the consensus sequence-[S/T]-X-C*, where * shows the end of the protein. B) Multiple sequence alignment of rat GlyT2 C-terminal region 791–799 from different species. Identical conserved PDZ binding motif (PBM) from different species are shown in bold. C) Pulldown experiment performed using three different baits: purified GST, GST fused to the C-terminus of GlyT2 or GST fused to the N-terminus of GlyT2. These were combined these with a cell lysate of COS7 cells expressing FLAG-tagged LNX1 isoform p80 (p80-LNX1) and amount of LNX1 interacting in each case was identified by immunoblotting against FLAG. D-E) COS7 cells expressing GlyT2 and FLAG-p80-LNX1 (D) or MYC-p70-LNX1 (E) were lysed and subjected to immunoprecipitation against GlyT2 and FLAG or MYC. Immunoprecipitates were analyzed by immunoblotting for indicated proteins, showing that GlyT2 coimmunoprecipitates both p70-LNX1 (E, middle lane) and p80-LNX1 (D, middle lane). Correspondingly, p70-LNX1 and p80-LNX1 coimmunoprecipitate GlyT2 (D, E, right lane). A higher exposure is shown in the case of p70 to observe GlyT2 in MYC-p70-LNX1 immunoprecipitates. F) Primary cortical neurons expressing FLAG-p80-LNX1 with or without GlyT2 were lysed and immunoprecipitated against GlyT2. Whereas no band for LNX1 was observed in GlyT2 immunoprecipitates when GlyT2 was not expressed, a clear band for FLAG-p80-LNX1 can be detected immunoprecipitating against GlyT2 in neurons expressing the transporter (F, right), indicating a specific interaction in neurons. G) COS7 cells expressing FLAG-p80-LNX1 and GlyT2 or GlyT2 with a deletion in the predicted interacting C-terminal region 791–799 (GlyT2-Δ8cter, fig 1B) were lysed and immunoprecipitated against GlyT2 and FLAG. GlyT2 immunoprecipitates present LNX1 in both cases, and correspondingly FLAG immunoprecipitates coimmunoprecipitate both variants of GlyT2. Note that intensity of the interaction is slightly lower in the case of GlyT2-Δ8cter using both direct and reverse immunoprecipitation.

We next explored the contribution of the PDZ binding motif (PBM) of GlyT2 to the global strength of the interaction by performing direct and reverse immunoprecipitation experiments as shown before. We repeated our experiments using a mutant of GlyT2 that lacks the last 8 amino acids found to bind the PDZ2 of LNX1^33^ (GlyT2-Δ8cter), which contains the conserved consensus PBM of GlyT2 (Fig 1B). The interaction between p80-LNX1 and GlyT2-Δ8cter seems slightly reduced, as signals of GlyT2 or LNX1 in both direct and reverse immunoprecipitations showed slightly lower intensity when compared to wild type GlyT2 experiments (Fig 1G). However, the interaction is still detectable (Fig 1G, right), suggesting that PDZ-PBM binding only contributes partially to the interaction. This result indicates that other domains of GlyT2 are necessary for the interaction and suggests the formation of additional interfaces during the PDZ-PBM interaction, which have been proposed to be critical to achieve affinity and specificity of modular domains such as PDZ or SH2/3^40–43^. In particular, it has been described PDZ-PBM interactions mediated by LNX1 require additional regions to achieve affinity in some cases, as occurs for example in the case of the LNX1-Numb interaction, which requires the internal phosphotyrosine binding domain of Numb^26^, or in the interaction of LNX1 with neuregulin-1/ErbB receptor and c-Src tyrosine kinase, which require additional regions in ErbB/c-Src C-terminal sequences^29,44^.

### RING finger domain of LNX1 ubiquitinates a C-terminal lysine cluster in GlyT2

LNX1 binds and mediates ubiquitination of several proteins including Numb^26^, ErbB2^44^, claudins^45^, CD8α^46^, PDZ-binding kinase^33^ or breakpoint cluster region protein (BCR)^33^. Ubiquitination of GlyT2 strongly modulates its total and surface expression, controlling its transport capacity^19^ and this control mechanism relays on the ubiquitination of a cytoplasmic C-terminal lysine cluster in GlyT2 (K751, K773, K787 and K791). Mutation of all four C-terminal lysines to arginines (GlyT2-4KR) results in reduced ubiquitination, impaired constitutive endocytosis and reduced PKC-mediated degradation of GlyT2^19^. We hypothesized that the LNX1-GlyT2 interaction may occur to control GlyT2 ubiquitination state, and therefore we first tested whether LNX1 ubiquitinates GlyT2 in cells. We coexpressed HA-tagged ubiquitin and GlyT2 with or without wild type p80-LNX1 in COS7 cells. We immunoprecipitated GlyT2 from cell lysates and measured its ubiquitination levels by immunoblotting with an anti-HA antibody, normalizing HA immunodetection against the immunoprecipitated amount of GlyT2 in each case to control for variability on GlyT2 expression. Compared to the control (Fig. 2A, lane 1), an increased ubiquitination signal was found when p80-LNX1 was overexpressed (Fig. 2A, lane 2) showing that increased levels of LNX1 indeed promote GlyT2 ubiquitination. These experiments also show that GlyT2 expression levels are reduced when p80-LNX1 is overexpressed (see next section). Quantification of these experiments indicated that overexpression of p80-LNX1 significantly increased GlyT2 ubiquitination levels by ~2 fold (Fig 1B, n=13, **p=0.004). To confirm that p80-LNX1 ubiquitin ligase activity is needed for this process, we inactivated p80-LNX1 ubiquitin ligase activity with a point mutation (C45A) that blunts RING finger ligase activity (p80-LNX1^C45A^)^26,30^. We coexpressed GlyT2 with inactive p80-LNX1^C45A^ and measured GlyT2 ubiquitination levels as before, finding that p80-LNX1^C45A^ failed to neither reduce GlyT2 expression levels or increase GlyT2 ubiquitination (Fig. 2A, lane 3; Fig 2B, n=13, p=0.99). Next, we explored whether this process requires the cytoplasmic C-terminal lysine cluster in GlyT2, as ubiquitination of these lysines have been previously demonstrated to control GlyT2 expression^19^. Using the same approach, we expressed the mutant GlyT2-4KR with or without p80-LNX1 or p80-LNX1^C45A^. Remarkably, we found no significant differences in ubiquitination levels of GlyT2-4KR when p80-LNX1 was overexpressed (Fig. 2A, lane 5; Fig 2B, n=5, p=1) and expression levels of GlyT2-4KR remained unaffected in these conditions. Also, as expected, p80-LNX1^C45A^ expression did not affect GlyT2-4KR ubiquitination (Fig. 2A, lane 6; Fig 2B, n=5, p=0.99). We found basal ubiquitination levels of GlyT2-4KR to be reduced as previously described^19^. As a control, we confirmed by immunoprecipitation that GlyT2-4KR maintains the ability of binding FLAG-p80-LNX1, demonstrating that the lack of ubiquitination in GlyT2-4KR is not a result of an impaired interaction (Fig. 2C). Taken together, these results indicate that p80-LNX1 induces ubiquitination of the C-terminal lysine cluster of GlyT2 through its functional RING finger domain.

**Figure 2.**
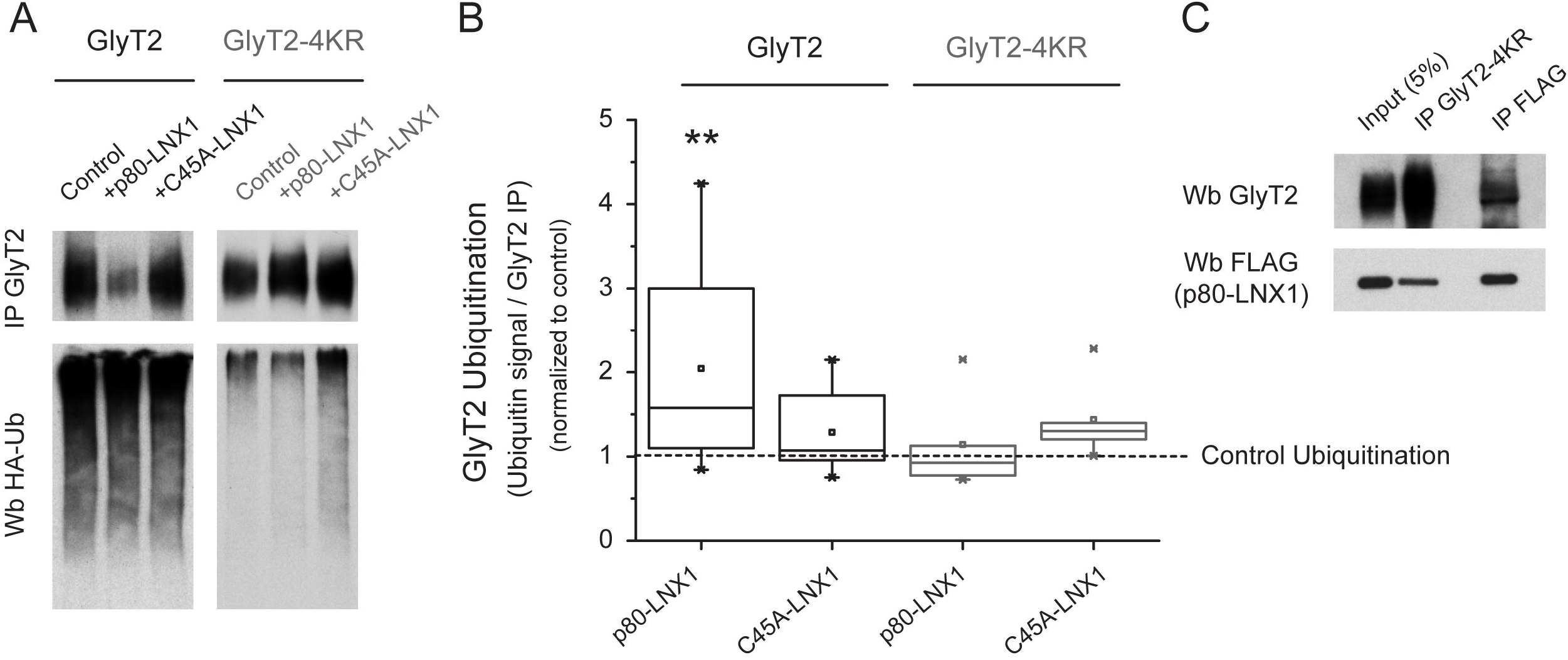
The RING-finger of LNX1 ubiquitinates the C-terminus lysine cluster of GlyT2. A) COS7 cells were transiently transfected with GlyT2 or GlyT2-4KR, HA-tagged ubiquitin and with or without wild type p80-LNX1 or functionally inactive p80-LNX1^C45A^ (showed in the figure as C45A-LNX1). GlyT2 was immunoprecipitated and ubiquitination of the transporter was assayed by immunoblotting against HA. Blots were probed against GlyT2 to normalize ubiquitination signal against the amount of GlyT2 immunoprecipitated in each case to correct for GlyT2 protein expression. B) Quantification of GlyT2 ubiquitination normalized to the control (no transfection of LNX1). Number of experiments in each case: p80-LNX1+GlyT2, n=13; C45A-LNX1+GlyT2, n = 13; p80-LNX1+GlyT2-4KR, n = 5; C45A-LNX1+GlyT2-4KR, n = 5. **p=0.004. C) COS7 cells expressing FLAG-p80-LNX1 and GlyT2-4KR were lysed and immunoprecipitated against GlyT2 and FLAG. GlyT2 immunoprecipitates present LNX1, and correspondingly FLAG immunoprecipitates coimmunoprecipitate GlyT2, indicating that GlyT2-4KR maintains the interaction with p80-LNX1.

### Ubiquitin ligase activity of LNX1 regulates GlyT2 expression

Our ubiquitination experiments did not find single bands corresponding to monoubiquitinated GlyT2 forms but showed a typical smear pattern that likely corresponds to polyubiquitinated forms^47,48^. Whereas monoubiquitination in general may regulate location and activity of diverse cellular proteins^49^, polyubiquitination is the type of ubiquitin modification that can target proteins for degradation^50^. Additionally, we noticed that in our ubiquitination experiments overexpression of p80-LNX1 induced a significant reduction in GlyT2 expression levels, suggesting that ubiquitination of GlyT2 by LNX1 regulates the expression of the transporter, similarly to what it does with other proteins^26,33,44–46^. To test this idea we coexpressed GlyT2 with p70-LNX1 (lacks RING finger, Fig. 1A) or wild type p80-LNX1 in COS7 cells and measured total and surface expression of GlyT2. Surface GlyT2 was measured using impermeable sulfo-NHS-SS-biotin to biotinylate and isolate surface proteins. Whereas coexpression with p70-LNX1 did not result in any significant change in GlyT2 expression (Fig 3A, C), overexpression of functionally active p80-LNX1 reduced GlyT2 both total and surface expression by ~65% and ~70%, respectively (Fig 3A, C, n=3). The ratio of biotinylated/total, however, remained unaffected in all cases, indicating that LNX1 impacts GlyT2 surface expression by regulating total expression and not by regulating surface levels *per se* (not shown). We confirmed the purity of surface protein isolation by tubulin immunoblotting, which as an intracellular protein does not undergo biotinylation (Fig 3A). The fact that p70 lacks ubiquitinating activity and fails to regulate GlyT2 expression suggests that ubiquitination is essential for this process. To confirm this, we coexpressed GlyT2 with inactive p80-LNX1^C45A^ and measured GlyT2 levels as before, finding that p80-LNX1^C45A^ failed to reduce GlyT2 expression levels (Fig. 3A, B). As a control, immunoprecipitation experiments confirmed that p80-LNX1^C45A^ still interacts with GlyT2 (data not shown), confirming that ubiquitin ligase activity of LNX1 regulates GlyT2 total expression, which results in a corresponding reduction of surface levels of the transporter.

**Figure 3.**
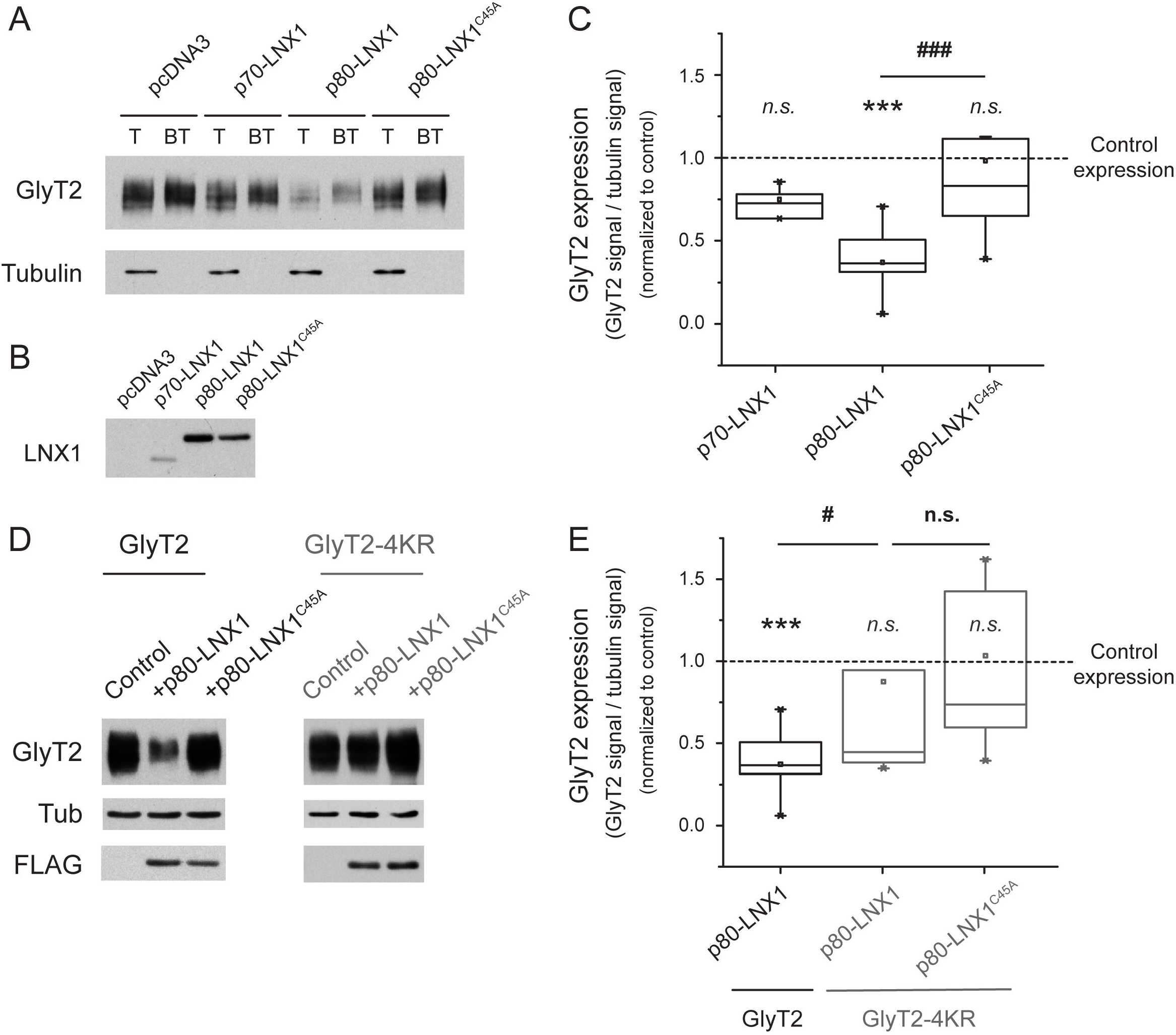
LNX1 controls GlyT2 expression. A) COS7 cells were transiently transfected with GlyT2 or GlyT2 and MYC-p70-LNX1, FLAG-p80-LNX1 or FLAG-p80-LNX1C45A and total and surface expression of GlyT2 was measured. Cell surface protein expression was isolated using impermeable sulfo-NHS-SS-biotin to biotinylate and isolate surface proteins using streptavidin agarose beads. T: input; BT: biotinylated. Tubulin is shown as a non-biotinylated intracellular control. B) Expression of LNX1 variants was confirmed by loading part of the input in a separate immunoblot. C) Quantification of the effect of LNX1 variants on GlyT2 expression. To correct for differential protein amount per lane, GlyT2 expression was normalized against tubulin, used as a loading control. GlyT2 expression in the different cases is shown normalized to the corrected amount of GlyT2 in the control with no LNX1 co-transfection (indicated by dashed line). *** p=2.6*10^−6^, ### p=1.6*10^−5^, using ANOVA with Tukey post hoc test. Quantification shown only for total expression of GlyT2 as surface fraction remained unaffected in all conditions. n(control)=10, n(p70)=4, n(p80)=10, n(C45A-p80)=10. D) COS7 cells were transiently transfected with GlyT2 or GlyT2-4KR and either FLAG-p80-LNX1 or FLAG-p80-LNX1C45A, and GlyT2 expression was measured by immunoblotting. Quantification of the effect of LNX1 variants on GlyT2 and GlyT2-4KR expression was normalized against tubulin as before. E) Quantification is shown normalized to the corrected signal in the control in each case with no LNX1 cotransfection (indicated by dashed line). *** p=2.6·10^−6^, # p=0.04, using ANOVA with Dunn-Sidak post hoc test. n(control)=10, n(p80)=10, n(GlyT2-4KR)=6, n(GlyT2-4KR-p80)=6, n(GlyT2-4KR-p80-LNX1^C45^)=6. Note that while p80-LNX1 induces a reduction of GlyT2 expression, GlyT2-4KR is highly protected in the same conditions. Effect of p80-LNX1 in wild type GlyT2 expression is shown here to ease comparison of the effects in GlyT2-4KR. FLAG immunobloting was used to confirm transfection of LNX1 variants.

LNX1 ubiquitinates the cytoplasmic C-terminal lysine cluster in GlyT2 (Fig 2A, B), which has been previously shown to modulate total and surface expression of the transporter^19^. Therefore, we reasoned that LNX1-mediated control of GlyT2 expression (Fig 3A–C) likely depends on the ubiquitination of the cytoplasmic C-terminal lysine cluster. To test this, we coexpressed wild-type GlyT2 or GlyT2-4KR together with p80-LNX1 or p80-LNX1^C48A^ and assayed GlyT2 expression levels as before. GlyT2 expression was reduced in the presence of p80-LNX1 as expected (Fig 3D, E; 37% of control, n=10; *** p= 8.9·10^−4^), but expression levels of GlyT2-4KR remarkably remained unaffected (~87% of control; n.s., p=0.9). Congruently, expression of p80-LNX1^C45A^ also did not change GlyT2-4KR levels (~103% of control; n.s., p=0.99). This results indicate that LNX1 can only control GlyT2 expression if the cytoplasmic C-terminal lysine cluster in GlyT2 remains intact, confirming that LNX1 promotes ubiquitination of those particular residues to control GlyT2 expression.

### GlyT2 and LNX1 colocalize in axons of primary neurons

GlyT2 is localized primarily in axons and terminals of glycinergic neurons^51^, where its activity is physiologically relevant for recapturing glycine to refill synaptic vesicles^1^. We next attempted to visualize the localization of LNX1 and GlyT2 in neurons. Our initial attempts of staining for LNX1 in spinal cord slices or primary cultures failed as available antibodies against LNX1 did not work well in these experiments and LNX proteins are generally difficult to detect since they are present at low levels in most adult tissues^28,29,44^ (see discussion). To overcome this, we transfected primary neurons with mCherry-tagged p80-LNX1 and GFP-GlyT2 and detected each protein by fluorescence microscopy using live cell imaging. Expression of mCherry-tagged p80-LNX1 together with GFP-GlyT2 showed expression of both proteins through all neuronal compartments, including axons, where they extensively co-localize (Fig 4A, B). p70-LNX1 expression was previously found presynaptically as well^31^, suggesting that both isoforms localize to similar regions in neurons. We confirmed that inactive mCherry-tagged p80-LNX1^C45A^ also co-localizes with GlyT2 in axons (Fig 4C), demonstrating that it fails to reduce GlyT2 expression levels (Fig 3A–D) because it lacks ubiquitin ligase activity, not because it does not localize where GlyT2 is expressed. Taken together, these experiments suggest that LNX1 can ubiquitinate and control protein expression of GlyT2 presynaptically, where GlyT2 transport of glycine is necessary for maintaining glycinergic neurotransmission strength.

**Figure 4.**
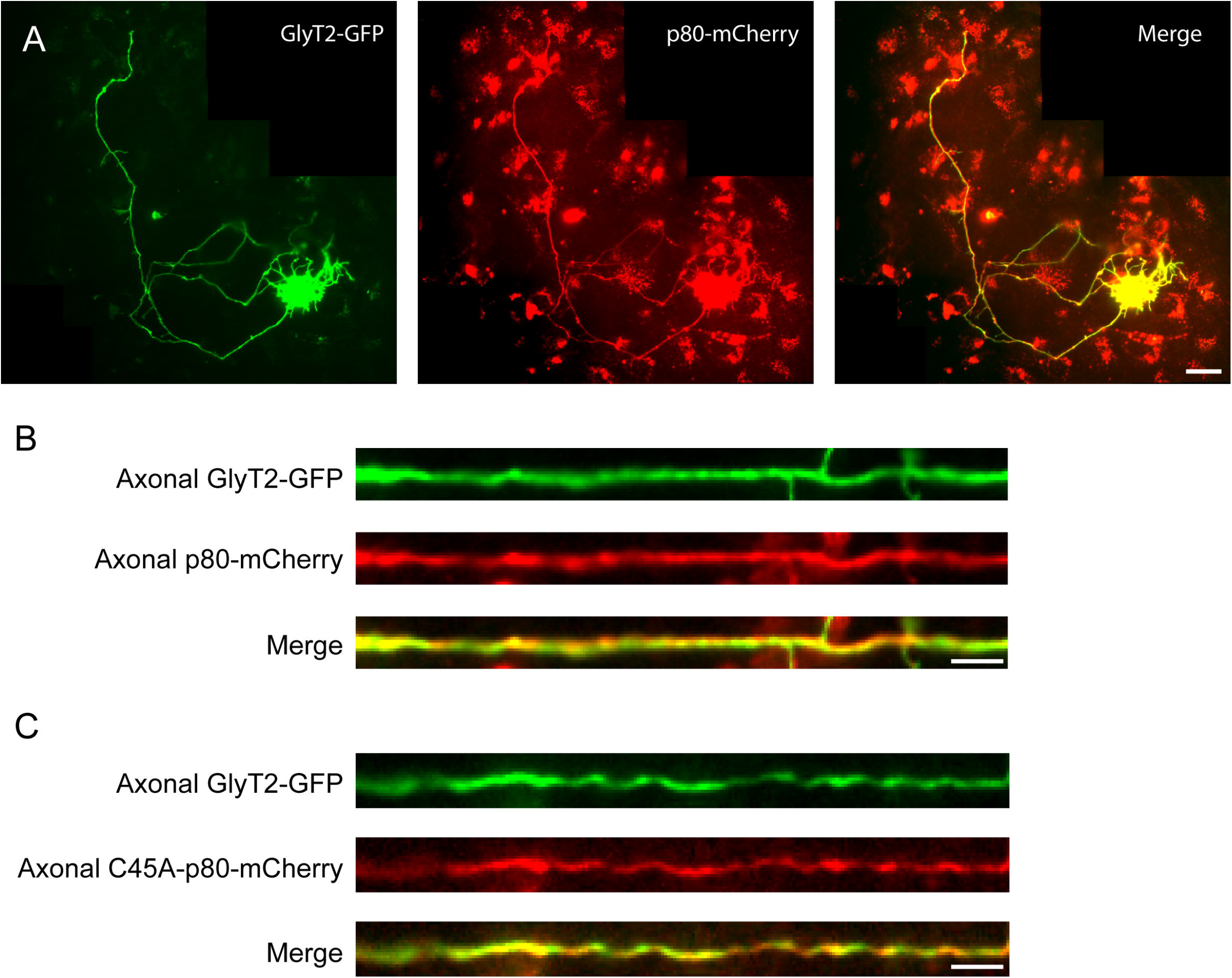
LNX1 colocalizes with GlyT2 in axons. A-C) Primary cortical neurons were transfected to express GFP-GlyT2 and mCherry-p80-LNX1 or mCherry-p80-LNX1^C45A^ and subjected to live cell microscopy at DIV9. Images of entire neurons are generated using the ImageJ stitching plugin. Simultaneous imaging of both GFP-GlyT2 and mCherry-LNX1 variants shows that both proteins present a similar localization in neurons. B-C) Higher magnification of axonal regions in neurons expressing mCherry-p80-LNX1 (B) or mCherry-p80-LNX1C45A (C).

### LNX1 controls GlyT2 transport activity

An increase in LNX1 expression results in a reduction of total and surface expression of GlyT2 (Fig 3). Glycine transport activity of GlyT2 depends on the availability of the transporter at the neuronal surface and, in fact, regulation of surface expression is a common regulatory mechanism in the solute carrier (SLC) family of transporters to control transport rates of a wide array of substrates across biological membranes of neurons^15,52–55^. Glycine transport by GlyT2 is essential for maintaining diverse aspects of motor function and impairment of its activity causes important pathologies in humans, from movement disorders to sensory dysfunctions^9^. To better understand the functional relevance of LNX1-mediated control of GlyT2 expression in transport activity, we next explored to what extent LNX1 expression may impact glycine recapture using [^3^H]-glycine uptake assays in COS7 cells and primary neurons. First, using COS7 cells we observed that coexpression of wild-type p80-LNX1 with GlyT2 resulted in a significant reduction of glycine transport of ~40% (Fig 5A, n(control)=43, n(p80-LNX1)=27; **** p=3.12*10^−10^). This effect, however, was not observed when GlyT2 was co-expressed with catalytically-inactive p80-LNX1^C45A^, in congruence with expression data (Fig 5A, n(control)=43, n(p80-LNX1^C45A^)=32; p=0.99). To test whether part of the effect on transport activity was partially due to a change in transport capacity of surface GlyT2 and not just a consequence of less availability of functional transporter molecules at the surface, we next tested the effect of p80-LNX1 on GlyT2-4KR, which is resistant to ubiquitination by p80-LNX1 (Fig 3). We transfected COS7 cells to express GlyT2-4KR and either p80-LNX1 or p80-LNX1^C45A^ and measured [^3^H]-glycine transport rates. Contrarily to wild type GlyT2, transport activity of GlyT2-4KR remained unaffected with p80-LNX1 overexpression (Fig 5A, right, n(GlyT2-4KR)=16, n(GlyT2-4KR, p80-LNX1)=16; p=0.69), indicating that p80-LNX1 does not alter the transport capacity of functional surface GlyT2 but does it by reducing the expression of the transporter. As expected, or p80-LNX1^C45A^ did not affect GlyT2-4KR transport activity (Fig 5A, right, n(GlyT2-4KR)=16, n(GlyT2-4KR, p80-LNX1^C45A^)=16; p=0.76). After dissecting the details of the molecular aspects of LNX1-mediated impact on GlyT2 glycine transport in COS7 cells, we next confirmed our main results in primary cultures of neurons. We transfected cortical neurons to express GlyT2 or GlyT2-4KR with or without p80-LNX1 and measured [^3^H]-glycine transport rates (see materials and methods). As expected, neuronal glycine transport by GlyT2 was strongly impacted by increased expression of p80-LNX1, resulting in a reduction of glycine uptake of ~55% (Fig 5B, n(control)=17, n(p80-LNX1)=18; ** p=0.01). Contrarily, GlyT2-4KR remained unaffected in the same conditions (Fig 5B, n(control)=17, n(p80-LNX1)=18; p=0.96), confirming that LNX1 can only control GlyT2 transport activity if the cytoplasmic C-terminal lysine cluster in GlyT2 remains intact. Additionally, we used transport assays in neurons to quantify the effect of p80-LNX1 on GlyT2-Δ8cter, as our immunoprecipitation experiments only showed a partial loss of the interaction in these conditions where the PDZ-PBM interaction is not possible (Fig 1G). We transfected neurons to express wild type GlyT2 or GlyT2-Δ8cter with p80-LNX1 and measured glycine transport rates. Single expression of GlyT2-Δ8cter in neurons showed a transport activity that was not significantly different from wild type GlyT2 (n(GlyT2)=17, n(GlyT2-Δ8cter)=10; p=0.99). However, when GlyT2-Δ8cter was coexpressed with p80-LNX1, glycine transport was not impacted, indicating that GlyT2-Δ8cter is protected against LNX1-mediated control of GlyT2 activity (Fig 5B, n(GlyT2-Δ8cter)=10, n(GlyT2-Δ8cter, p80-LNX1)=10; p=0.94). These experiments confirm that PDZ-PBM binding contributes to the interaction between GlyT2 and LNX1 as suggested by immunoprecipitation assays (Fig 1G), and confirm that this region of GlyT2 is essential for the control exerted by LNX1 on GlyT2 activity in neurons.

**Figure 5.**
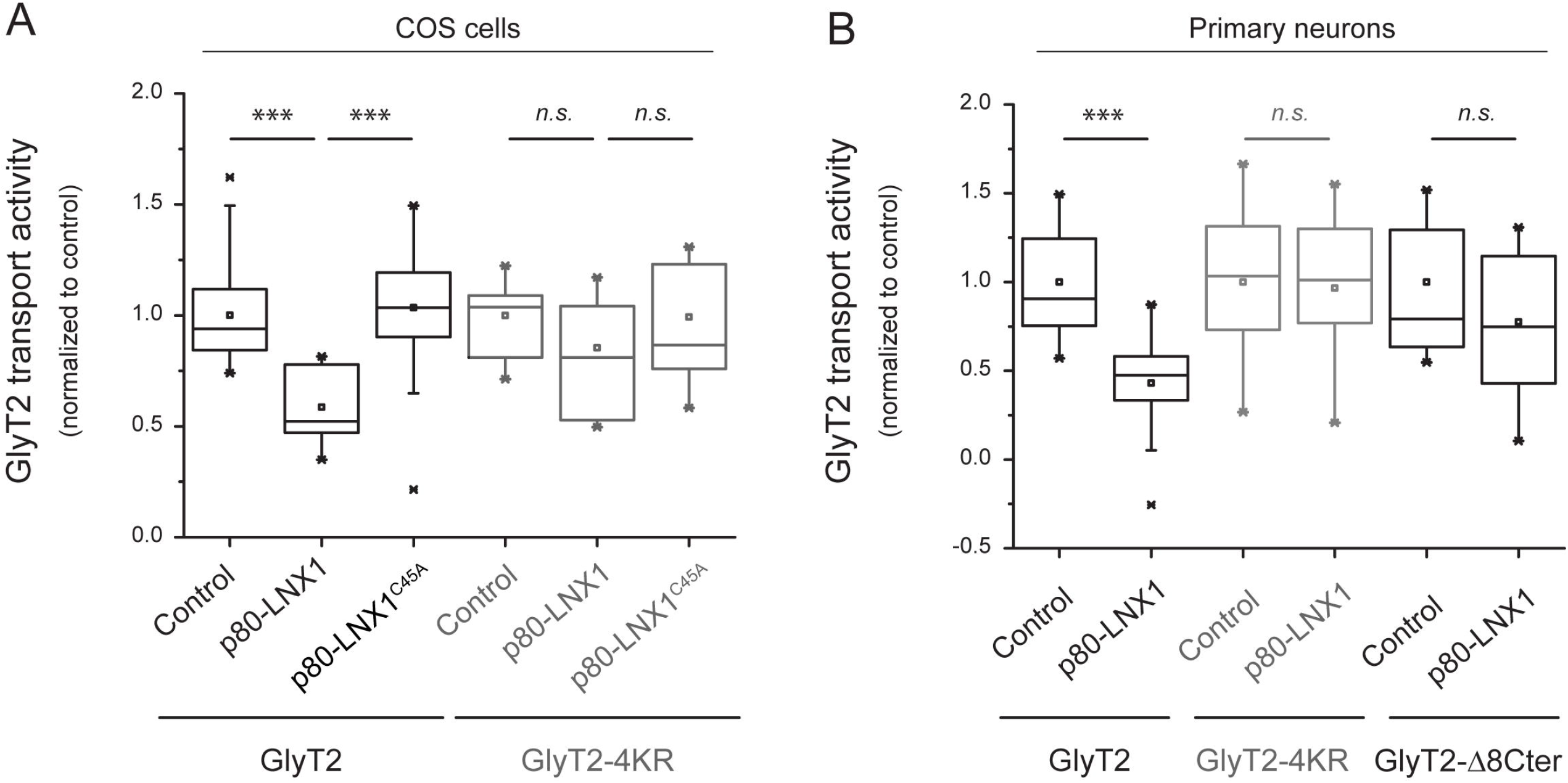
LNX1 ubiquitin ligase activity controls glycine recapture by GlyT2. A-B) COS7 cells were transiently transfected with GlyT2 or GlyT2-4KR in combination with pcDNA3, p80-LNX1 or p80-LNX1^C45A^ and glycine transport rates were measured using [3H]-Glycine transport assays as described in materials and methods. Glycine transport is shown normalized against control conditions in each case, where the actual values for transport rates were (in pmol of Gly/mg of protein/min): GlyT2: 201.7 ± 6.3; GlyT2-4KR: 363.6 ± 15.0; C-D) Primary cortical neurons were transfected to express either GlyT2, GlyT2-4KR or GlyT2-Δ8cter in combination with pcDNA3or p80-LNX1. Glycine transport rates were measured using [3H]-Glycine transport assays. Glycine transport is shown normalized against control conditions in each case, where the actual values for transport rates were (in pmol of Gly/mg of protein/min): GlyT2: 11.3 ± 0.9; GlyT2-4KR: 17.1 ± 1.6; GlyT2-Δ8cter: 12.8 ± 1.5; control, p < 0.001; n.s., not significantly different. Comparisons of the means were performed using ANOVA by Tukey’s post hoc test.

## DISCUSSION

This study provides compelling novel evidence that demonstrates that the RING-finger E3 ubiquitin ligase LNX1 is a functional regulator of the neuronal glycine transporter GlyT2. Our data using interaction assays, imaging and glycine transport assays reveal that LNX1 interacts with the C-terminus of GlyT2 in neurons and ubiquitinates a C-terminal cluster of lysines of this transporter to control its expression and activity.

LNX1 (Ligand of Numb Protein-X) or PDZRN (PDZ and RING) is part of the LNX family of proteins, which consists of five members (LNX1-5) that present a structure containing both an n-terminus RING-finger domain and one to four PDZ domains in tandem in the C-terminus^56–58^. This modular domain organization, unique to the LNX family, suggests that LNX members have a facility for ubiquitinating substrates containing protein binding motifs (PBM), which are recognized specifically through the different PDZ domains. LNX1 has been shown to localize to axons and presynaptic terminals^31^ and it interacts with a set of presynaptic proteins including CAST^31^, PKCα^30,59^, ERC1, ERC2 and LIPRIN-αs^32^. LNX1 also interacts with NUMB^26,60^ and c-Src^29^, which are also expressed in presynaptic terminals^61,62^, and it can ubiquitinate these substrates to target them for degradation^26,29,30^. Our work demonstrates a similar functional relationship between LNX1 and the presynaptic glycine transporter GlyT2, as we show that LNX1-mediated ubiquitination of GlyT2 controls its expression and activity.

Given the tandem PDZ organization of the LNX family, it has been proposed that LNX proteins may act as molecular scaffolds to localize their interacting partners to the same specific subcellular location^30,56^. In neurons, LNX1 may act as a presynaptic scaffold, promoting the formation of multimolecular complexes that could allow the correct subcellular localization of GlyT2 and other interacting partners^31,63^. GlyT2 has been previously shown to interact with several presynaptic proteins, including syntaxin1^16^, Plasma Membrane Calcium ATPases PMCA2 and PMC A3^18^ and sodium/potassium ATPase subunits α3 (α3NKA) and β2 (β2NKA)^17^. It is possible that LNX1 may act as a scaffold to facilitate the interaction of GlyT2 with the mentioned proteins, and this idea of a GlyT2-LNX1-PMCA/NKA multimolecular complex is supported by a recent proteomic study that found PMCA2 and β2NKA as candidate interacting partners of LNX1^32^.

LNX1 and the very close isoform LNX2 are expressed in many adult tissues^63^ and a recent study has reported that in the spinal cord the mRNAs of these isoforms are primarily found in neurons^28^. Correspondingly, GlyT2 is expressed as well in neurons the spinal cord, where it is considered a marker for glycinergic neurons^51,64^. However and despite the fact that LNXs mRNAs are easily detected^28,60,63^, LNX proteins are present at low levels in most adult tissues^28,29,44^. This suggests that either LNX mRNAs do not undergo significant translation or that LNX proteins present a very short half-life. Our results support the second idea, as we found that the half-life of transfected p80-LNX1 in primary neurons is remarkably short, observing full degradation of the overexpressed protein after 48h while virtually every other protein tested, including GlyT2, did not change their expression levels in these conditions (data not shown; see methods). This suggests that the function of LNX proteins might be regulated temporarily through the control of their protein expression, which at steady state would be maintained at low levels but could be rapidly increased when an increase in their activity is needed. This mechanism would not be unique to LNX proteins, as it has been described that increased translation of existing transcripts can help to quickly synthesize newly required proteins that control short-term temporal adaptation mechanisms^65^. Even more, E3-substrate interactions are in many cases constitutive, and, in such cases, regulation is thought to occur at the level of E3 transcription/translation or degradation^66^. This paradigm could be particularly relevant for neuronal function, as ubiquitination has been shown to acutely regulate neurotransmitter release in mammalian neurons^67^.

Ubiquitination is major pathway that controls turnover of neuronal membrane proteins^22^. In particular, ubiquitination strongly regulates GlyT2 surface expression, as ubiquitination of the C-terminus lysine cluster of GlyT2 is required for constitutive endocytosis and sorting into the recycling pathway^19^. Additionally, GlyT2 ubiquitination status in neurons is highly responsive to the free pool of ubiquitin, which when pharmacologically perturbed results in altered GlyT2 turnover and surface expression^19^. Although ubiquitination is a major regulatory pathway that controls GlyT2 function, the molecular identity of the enzymes catalyzing GlyT2 ubiquitination has remained elusive. Our work presented here identifies for the first time an E3-ubiquitin ligase acting on GlyT2. We discovered that the E3 ubiquitin ligase LNX1 modulates the ubiquitination status of GlyT2, ubiquitinating a cytoplasmic C-terminal lysine cluster in GlyT2 (K751, K773, K787 and K791) in a process that regulates the expression levels and transport activity of GlyT2 in neurons. Our experiments show that both proteins co-localize in axons, where GlyT2 transport is needed. Correct maintenance of glycine transport by GlyT2 is essential for human physiology, as GlyT2 is an essential regulator of glycinergic neurotransmission strength and therefore the new regulatory mechanism discovered here may have pathophysiological relevance on the biology of glycinergic neurotransmission and might help frame future investigations into the molecular basis of human disease states that are consequence of presynaptic glycinergic neurotransmission dysfunction.

## Bibliography

1. Apostolides, P. F. & Trussell, L. O. Rapid, activity-independent turnover of vesicular transmitter content at a mixed glycine/GABA synapse. J. Neurosci. 33, 4768–81 (2013).

2. Gomeza, J. et al. Deletion of the Mouse Glycine Transporter 2 Results in a Hyperekplexia Phenotype and Postnatal Lethality. Neuron 40, 797–806 (2003).

3. Suhren, O., Bruyn G W & Tuynman J A. Hyperexplexia: A hereditary startle syndrome. J. Neurol. Sci. 3, 577–605 (1966).

4. Harvey, R. J., Topf, M., Harvey, K. & Rees, M. I. The genetics of hyperekplexia: more than startle! Trends in Genetics 24, 439–447 (Elsevier Current Trends, 2008).

5. Schaefer, N., Langlhofer, G., Kluck, C. J. & Villmann, C. Glycine receptor mouse mutants: model systems for human hyperekplexia. Br. J. Pharmacol. 170, 933–952 (2013).

6. Carta, E. et al. Mutations in the GlyT2 gene (SLC6A5) are a second major cause of startle disease. J. Biol. Chem. 287, 28975–85 (2012).

7. Lynch, J. W. & Callister, R. J. Glycine receptors: a new therapeutic target in pain pathways. Curr. Opin. Investig. Drugs 7, 48–53 (2006).

8. Grothe, B. Sensory systems: New roles for synaptic inhibition in sound localization. Nat. Rev. Neurosci. 4, 540–550 (2003).

9. Zafra, F., Ibáñez, I. & Giménez, C. Glycinergic transmission: glycine transporter GlyT2 in neuronal pathologies. Neuronal Signal. 1, NS20160009 (2016).

10. Fornes, A. et al. Trafficking properties and activity regulation of the neuronal glycine transporter GLYT2 by protein kinase C. Biochem J 412, 495–506 (2008).

11. de Juan-Sanz, J., Zafra, F., López-Corcuera, B. & Aragón, C. Endocytosis of the neuronal glycine transporter GLYT2: role of membrane rafts and protein kinase C-dependent ubiquitination. Traffic 12, 1850–67 (2011).

12. Arribas-González, E., de Juan-Sanz, J., Aragón, C. & López-Corcuera, B. Molecular basis of the dominant negative effect of a glycine transporter 2 mutation associated with hyperekplexia. J. Biol. Chem. 290, 2150–65 (2015).

13. Arribas-González, E., Alonso-Torres, P., Aragón, C. & López-Corcuera, B. Calnexin-assisted biogenesis of the neuronal glycine transporter 2 (GlyT2). PLoS One 8, e63230 (2013).

14. Jiménez, E. et al. P2Y purinergic regulation of the glycine neurotransmitter transporters. J. Biol. Chem. 286, 10712–24 (2011).

15. Villarejo-López, L. et al. P2X receptors up-regulate the cell-surface expression of the neuronal glycine transporter GlyT2. Neuropharmacology 125, 99–116 (2017).

16. Geerlings, A., Núñez, E., López-Corcuera, B. & Aragón, C. Calcium- and syntaxin 1-mediated trafficking of the neuronal glycine transporter GLYT2. J. Biol. Chem. 276, 17584–90 (2001).

17. de Juan-Sanz, J. et al. Na+/K+-ATPase is a new interacting partner for the neuronal glycine transporter GlyT2 that downregulates its expression in vitro and in vivo. J. Neurosci. 33, 14269–81 (2013).

18. de Juan-Sanz, J. et al. Presynaptic control of glycine transporter 2 (GlyT2) by physical and functional association with plasma membrane Ca2+-ATPase (PMCA) and Na+-Ca2+ exchanger (NCX). J. Biol. Chem. M114.586966-(2014). doi:10.1074/jbc.M114.586966

19. de Juan-Sanz, J. et al. Constitutive endocytosis and turnover of the neuronal glycine transporter GlyT2 is dependent on ubiquitination of a C-terminal lysine cluster. PLoS One 8, e58863 (2013).

20. DiAntonio, A. et al. Ubiquitination-dependent mechanisms regulate synaptic growth and function. Nature 412, 449–452 (2001).

21. Pinto, M. J. et al. The proteasome controls presynaptic differentiation through modulation of an on-site pool of polyubiquitinated conjugates. J. Cell Biol. 212, (2016).

22. Schwarz, L. A. & Patrick, G. N. Ubiquitin-dependent endocytosis, trafficking and turnover of neuronal membrane proteins. Mol. Cell. Neurosci. 49, 387–393 (2012).

23. Berndsen, C. E. & Wolberger, C. New insights into ubiquitin E3 ligase mechanism. Nat. Struct. Mol. Biol. 21, 301–307 (2014).

24. Nakayama, K. I. & Nakayama, K. Ubiquitin ligases: cell-cycle control and cancer. Nat. Rev. Cancer 6, 369–381 (2006).

25. Deshaies, R. J. & Joazeiro, C. A. P. RING Domain E3 Ubiquitin Ligases. Annu. Rev. Biochem. 78, 399–434 (2009).

26. Nie, J. et al. LNX functions as a RING type E3 ubiquitin ligase that targets the cell fate determinant Numb for ubiquitin-dependent degradation. EMBO J. 21, 93–102 (2002).

27. Gulino, A., Di Marcotullio, L. & Screpanti, I. The multiple functions of Numb. Exp. Cell Res. 316, 900–6 (2010).

28. Lenihan, J. A., Saha, O., Mansfield, L. M. & Young, P. W. Tight, cell type-specific control of LNX expression in the nervous system, at the level of transcription, translation and protein stability. Gene 552, 39–50 (2014).

29. Weiss, A., Baumgartner, M., Radziwill, G., Dennler, J. & Moelling, K. c-Src is a PDZ interaction partner and substrate of the E3 ubiquitin ligase Ligand-of-Numb protein X1. FEBS Letters 581, (2007).

30. Wolting, C. D. et al. Biochemical and Computational Analysis Of LNX1 Interacting Proteins. PLoS One 6, e26248 (2011).

31. Higa, S., Tokoro, T., Inoue, E., Kitajima, I. & Ohtsuka, T. The active zone protein CAST directly associates with Ligand-of-Numb protein X. Biochem. Biophys. Res. Commun. 354, 686–692 (2007).

32. Lenihan, J. A. et al. Decreased Anxiety-Related Behaviour but Apparently Unperturbed NUMB Function in Ligand of NUMB Protein-X (LNX) 1/2 Double Knockout Mice. Mol. Neurobiol. (2016). doi:10.1007/s12035-016-0261-0

33. Guo, Z. et al. Proteomics strategy to identify substrates of LNX, a PDZ domain-containing E3 ubiquitin ligase. J. Proteome Res. 11, 4847–62 (2012).

34. Song, E. et al. A high efficiency strategy for binding property characterization of peptide-binding domains. Mol. Cell. Proteomics 5, 1368–81 (2006).

35. Zafra, F. et al. Glycine transporters are differentially expressed among CNS cells. J. Neurosci. 15, 3952–3969 (1995).

36. Núñez, E. et al. Subcellular localization of the neuronal glycine transporter GLYT2 in brainstem. Traffic 10, 829–43 (2009).

37. Giménez, C. et al. A novel dominant hyperekplexia mutation Y705C alters trafficking and biochemical properties of the presynaptic glycine transporter GlyT2. J. Biol. Chem. 287, 28986–9002 (2012).

38. de Juan-Sanz, J. et al. Axonal Endoplasmic Reticulum Ca 2+ Content Controls Release Probability in CNS Nerve Terminals. Neuron 93, 867–881.e6 (2017).

39. Sambrook, J. & Russell, D. W. Detection of Protein-Protein Interactions Using the GST Fusion Protein Pulldown Technique. Cold Spring Harb. Protoc. 2006, pdb.prot3757 (2006).

40. Bae, J. H. et al. The Selectivity of Receptor Tyrosine Kinase Signaling Is Controlled by a Secondary SH2 Domain Binding Site. Cell 138, 514–524 (2009).

41. Lee, C. H., Saksela, K., Mirza, U. A., Chait, B. T. & Kuriyan, J. Crystal structure of the conserved core of HIV-1 Nef complexed with a Src family SH3 domain. Cell 85, 931–42 (1996).

42. Ostermeier, C. & Brunger, A. T. Structural basis of Rab effector specificity: crystal structure of the small G protein Rab3A complexed with the effector domain of rabphilin-3A. Cell 96, 363–74 (1999).

43. Pascoe, H. G. et al. Secondary PDZ domain-binding site on class B plexins enhances the affinity for PDZ–RhoGEF. Proc. Natl. Acad. Sci. 112, 14852–14857 (2015).

44. Young, P. et al. LNX1 is a perisynaptic Schwann cell specific E3 ubiquitin ligase that interacts with ErbB2. Mol. Cell. Neurosci. 30, 238–248 (2005).

45. Takahashi, S. et al. The E3 ubiquitin ligase LNX1p80 promotes the removal of claudins from tight junctions in MDCK cells. J. Cell Sci. 122, 985–994 (2009).

46. D’Agostino, M. et al. Ligand of Numb proteins LNX1p80 and LNX2 interact with the human glycoprotein CD8 and promote its ubiquitylation and endocytosis. J. Cell Sci. 124, 3545–3556 (2011).

47. Xie, P. et al. The covalent modifier Nedd8 is critical for the activation of Smurf1 ubiquitin ligase in tumorigenesis. Nat. Commun. 5, 3733 (2014).

48. Nguyen, A. T. et al. UBE2O remodels the proteome during terminal erythroid differentiation. Science 357, eaan0218 (2017).

49. Hicke, L. Protein regulation by monoubiquitin. Nat. Rev. Mol. Cell Biol. 2, 195–201 (2001).

50. Weissman, A. M. Themes and variations on ubiquitylation. Nat. Rev. Mol. Cell Biol. 2, 169–178 (2001).

51. Poyatos, I., Ponce, J., Aragón, C., Giménez, C. & Zafra, F. The glycine transporter GLYT2 is a reliable marker for glycine-immunoreactive neurons. Brain Res. Mol. Brain Res. 49, 63–70 (1997).

52. Ashrafi, G., Wu, Z., Farrell, R. J. & Ryan, T. A. GLUT4 Mobilization Supports Energetic Demands of Active Synapses. Neuron 93, 606–615.e3 (2017).

53. Richardson, B. D. et al. Membrane potential shapes regulation of dopamine transporter trafficking at the plasma membrane. Nat. Commun. 7, 10423 (2016).

54. García-Tardón, N. et al. Protein Kinase C (PKC)-promoted Endocytosis of Glutamate Transporter GLT-1 Requires Ubiquitin Ligase Nedd4-2-dependent Ubiquitination but Not Phosphorylation. J. Biol. Chem. 287, 19177–19187 (2012).

55. Lamothe, S. M. & Zhang, S. Chapter Five - Ubiquitination of Ion Channels and Transporters. Prog. Mol. Biol. Transl. Sci. 141, 161–223 (2016).

56. Flynn, M., Saha, O. & Young, P. Molecular evolution of the LNX gene family. BMC Evol. Biol. 11, 235 (2011).

57. Zheng, N., Wang, P., Jeffrey, P. D. & Pavletich, N. P. Structure of a c-Cbl-UbcH7 complex: RING domain function in ubiquitin-protein ligases. Cell 102, 533–9 (2000).

58. Lorick, K. L. et al. RING fingers mediate ubiquitin-conjugating enzyme (E2)-dependent ubiquitination. Proc. Natl. Acad. Sci. U. S. A. 96, 11364–9 (1999).

59. Fioravante, D. et al. Protein kinase C is a calcium sensor for presynaptic short-term plasticity. Elife 3, (2014).

60. Dho, S. E. et al. The mammalian numb phosphotyrosine-binding domain. Characterization of binding specificity and identification of a novel PDZ domain-containing numb binding protein, LNX. J. Biol. Chem. 273, 9179–87 (1998).

61. Nishimura, T. et al. CRMP-2 regulates polarized Numb-mediated endocytosis for axon growth. Nat. Cell Biol. 5, 819–26 (2003).

62. Onofri, F. et al. Synapsin I interacts with c-Src and stimulates its tyrosine kinase activity. Proc. Natl. Acad. Sci. U. S. A. 94, 12168–73 (1997).

63. Rice, D. S., Northcutt, G. M. & Kurschner, C. The Lnx family proteins function as molecular scaffolds for Numb family proteins. Mol. Cell. Neurosci. 18, 525–40 (2001).

64. Zeilhofer, H. U. The glycinergic control of spinal pain processing. Cell. Mol. Life Sci. 62, 2027–35 (2005).

65. Liu, Y., Beyer, A. & Aebersold, R. On the Dependency of Cellular Protein Levels on mRNA Abundance. Cell 165, 535–550 (2016).

66. Metzger, M. B., Pruneda, J. N., Klevit, R. E. & Weissman, A. M. RING-type E3 ligases: master manipulators of E2 ubiquitin-conjugating enzymes and ubiquitination. Biochim. Biophys. Acta 1843, 47–60 (2014).

67. Rinetti, G. V. & Schweizer, F. E. Ubiquitination Acutely Regulates Presynaptic Neurotransmitter Release in Mammalian Neurons. J. Neurosci. 30, 3157–3166 (2010).

